# Structural and biochemical insights on the mechanism of action of the clinical USP1 inhibitor, KSQ-4279

**DOI:** 10.1101/2024.05.16.594330

**Authors:** ML Rennie, M Gundogdu, C Arkinson, S Liness, S Frame, H Walden

## Abstract

DNA damage triggers cell signalling cascades that mediate repair. This signalling is frequently dysregulated in cancers. The proteins that mediate this signalling are potential targets for therapeutic intervention. Ubiquitin-specific protease 1 (USP1) is one such target, with small molecule inhibitors already in clinical trials. Here we use biochemical assays and cryo-electron microscopy (cryo-EM) to study the clinical USP1 inhibitor, KSQ-4279 (RO7623066), and compare this to the well-established, tool compound, ML323. We find that KSQ-4279 binds to the same cryptic site of USP1 as ML323 but disrupts the protein structure in subtly different ways. Inhibitor binding drives a substantial increase in thermal stability of USP1, which may be mediated through the inhibitors filling a hydrophobic tunnel in USP1. Our results contribute to the understanding of the mechanism of action of USP1 inhibitors at the molecular level.

## Introduction

Ubiquitin-specific protease 1 (USP1) deubiquitinates substrates involved in DNA repair. It functions with a cofactor protein, USP1-associated factor 1 (UAF1), that stimulates enzymatic activity and assists substrate engagement^1–3^. These substrates include the DNA clamps – PCNA and FANCI-FANCD2, involved in translesion synthesis^4,5^ and the Fanconi Anemia pathway^6–10^, respectively. Mono-ubiquitination of PCNA recruits polymerases that can bypass sites of DNA damage^11^, while mono-ubiquitination of FANCI-FANCD2 appears to lock the complex onto chromatin^7–10,12,13^. Deubiquitination by USP1 is presumed to reverse these processes to ensure genomic integrity during DNA repair.

USP1 is emerging as a potential target in the treatment of several cancers. It has long been postulated that synthetic lethality – simultaneous disruption of two or more non-essential genes/proteins resulting in cell death – may yield targeted cancer therapies^14^. This is exemplified by poly(ADP-ribose) polymerase (PARP) inhibitors for the treatment of *BRCA1/2*-mutant tumours^15,16^. However, tumours may develop resistance through a number of mechanisms^17^. The *USP1* gene is upregulated in several types of cancer and this often correlates with poor prognosis^18–22^, with some of these tumours also containing mutations in the *BRCA1* gene^19^. Synthetic lethality between *USP1* and *BRCA1/2* has been demonstrated with CRISPR/CAS9 screens and USP1 inhibitors^23,24^. The combination of PARP inhibitors and USP1 inhibitors in *BRCA1/2*-mutant tumours is even more effective than PARP inhibitors or USP1 inhibitors alone in this genetic sub-population^23,24^ and may provide a means for overcoming PARP inhibitor resistance. There are currently several USP1 inhibitors in phase 1 clinical trials – KSQ-4279/RO7623066 (KSQ Therapeutics/Hoffmann-La Roche), TNG348 (Tango Therapeutics), ISM3091/XL309 (In Silico Medicine/Exelixis), SIM0501 (Simcere Jiangsu Pharmaceutical Co), and HSK39775 (Xizang Haisco Pharmaceutical Co). Mechanistically, PARP inhibitors block enzymatic activity and trap PARP1 on DNA^25^; in fact PARP1 inhibition is more cytotoxic than PARP1 removal^25^. Curiously, there is evidence that USP1 can also become trapped on DNA when its enzymatic activity is impaired^26^. However, disruption of PCNA homeostasis appears to be critical to the cytotoxicity of USP1 inhibition^24^.

The first selective USP1 inhibitor, ML323, was developed by the Zhuang and Maloney groups^27,28^. We recently determined the structure of this inhibitor in complex with USP1, UAF1, and the FANCI-FANCD2^Ub^ substrate using cryo-electron microscopy (cryo-EM)^29^. USP1 is inhibited by ML323 via interaction with a cryptic binding site, situated between the palm and thumb subdomains of the USP fold, that is almost completely obscured in the absence of inhibitor^29^ (PDB IDs: 7AY0, 7AY2^3^, 7ZH3^29^). We sought to elucidate the binding mode of the clinical USP1 inhibitor, KSQ-4279, and compare this to ML323. We found that both ML323 and KSQ-4279 drive a substantial increase in stability of the USP fold of USP1. Cryo-EM revealed a similar binding mode between the two compounds, with both compounds occupying a hydrophobic tunnel. KSQ-4279 drives subtle rearrangements in the cryptic site compared with ML323 and the compounds result in different levels of disorder in the adjacent regions. Our data are consistent with an induced fit binding to USP1 for both inhibitors.

## Results and Discussion

### Biochemical comparison

We first compared ML323 and KSQ-4279 inhibition in ubiquitin-rhodamine assays using the DUB*profiler*™ assay (Ubiquigent). Screening against a panel of almost 50 deubiquitinase enzymes revealed that both ML323 and KSQ-4279 were selective against USP1 at 0.01 μM of inhibitor (Figure 1A). KSQ-4279 retained exquisite selectivity for USP1 at inhibitor concentrations as high as 10,000 times the IC_50_ value (data not shown). In contrast, ML323 showed inhibition of USP12 and USP46 at concentrations 100 times higher than the IC_50_ value for USP1. USP12 and USP46 are two close homologs of USP1 that also bind UAF1^30–33^. Both ML323 and KSQ-4279 resulted in near complete inhibition of USP1-UAF1.

**Figure 1:**
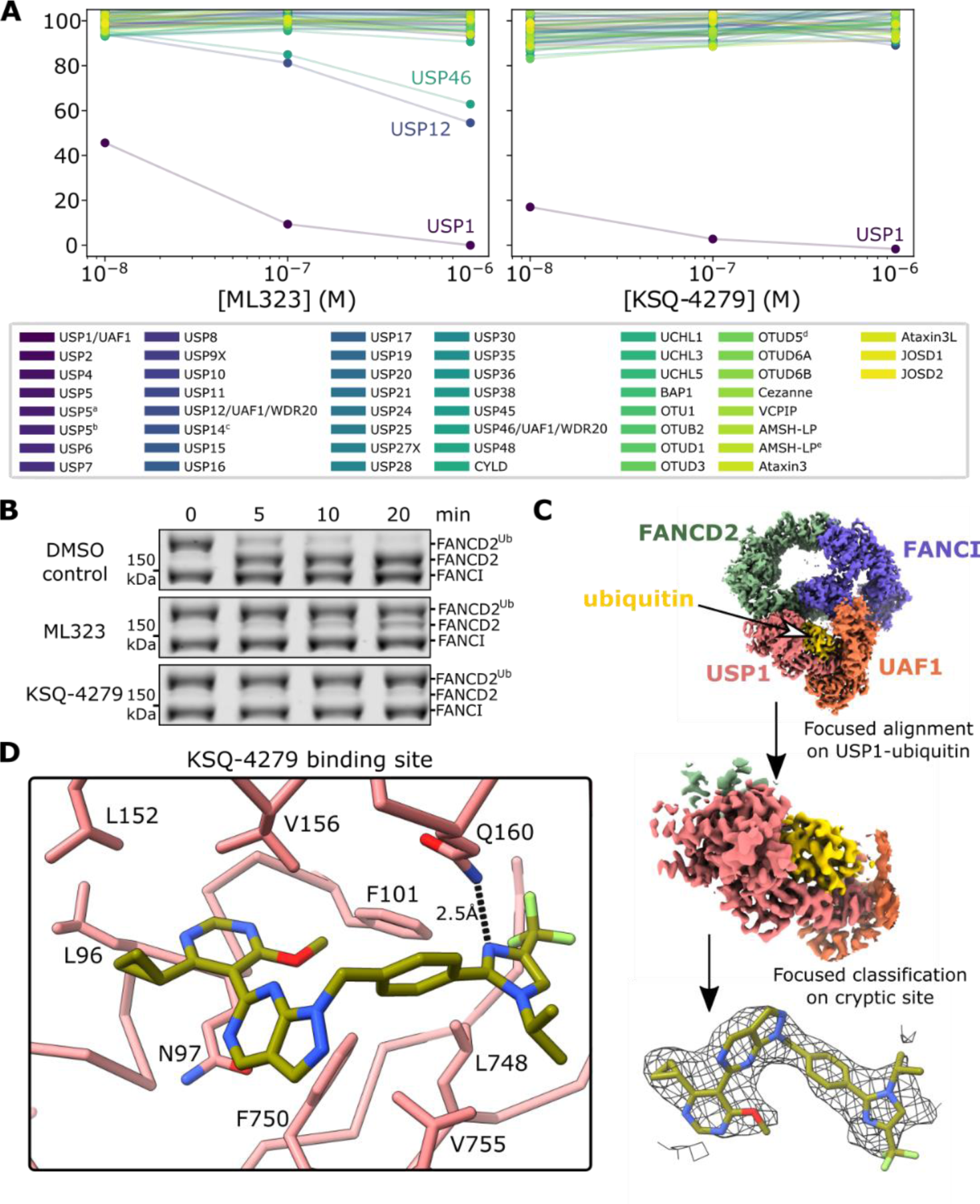
Biochemical and structural characterization of KSQ-4279. (A) Evaluation of the selectivity of ML323 and KSQ-4279 inhibitors across the DUBprofiler™ panel (Ubiquigent). ^a^enzyme activated half-maximally with additional ubiquitin; ^b^enzyme activated maximally with additional ubiquitin; ^c^enzyme activated with proteasome-V5; ^d^pS177; ^e^with additional zinc added. (B) Gel-based assay to demonstrate USP1 activity against FANCI-FANCD2^Ub^-dsDNA in the presence or absence of ML323 or KSQ-4279 (25 µM). USP1-UAF1 enzyme and FANCI-FANCD2^Ub^ substrate were used at 0.01 µM and 1 µM, respectively. Two technical replicates were performed. (C) Cryo-EM analysis of USP1^C90S^-UAF1-FANCI-FANCD2^Ub^-dsDNA with KSQ-4279. Density within 2.5 Å of KSQ-4279 is shown at 7.1σ (threshold of 0.085; sharpened map). (D) KSQ-4279 binding site. Sidechains of residues contacting KSQ-4279 and protein Cα atoms are shown; black dashed lines indicate hydrogen bonds.

### Structural characterization of KSQ-4279 binding

In order to determine the structure of KSQ-4279 bound to USP1 we employed the FANCI-FANCD2^Ub^ substrate to trap a larger complex more amenable to cryo-electron microscopy, similar to our approach with ML323^29^. Excess KSQ-4279 reduced the extent of deubiquitination of FANCI-FANCD2^Ub^-dsDNA substrate in reconstitution, gel-based assays (Figure 1B). After 20 minutes incubation with enzyme almost no deubiquitination was observed in the presence of KSQ-4279, contrasting with results obtained in the presence of excess ML323 for which deubiquitinated FANCD2 was apparent. This suggests KSQ-4279 disrupts FANCI-FANCD2^Ub^-dsDNA deubiquitination to a greater extent than ML323. Cryo-EM analysis of the C90S active site mutant of USP1 reconstituted with its cofactor UAF1 and FANCI-FANCD2^Ub^-dsDNA substrate, in the presence of KSQ-4279, allowed reconstruction of USP1-ubiquitin with this inhibitor at a resolution of ∼3.2 Å (Figure 1C, Figure S1, Table S1). The datasets for both inhibitor-bound and inhibitor-free particles and 3D classification was used to identify a subset consistent with inhibitor-bound USP1, similarly to ML323^29^ (Figure S1). KSQ-4279 binds the same cryptic pocket as ML323, between the palm and thumb subdomains, replacing several residues of the hydrophobic core (Figure 1D). We refer to these residues, 83-88, as the Replaced by Inhibitor Region (RIR). The conformation of KSQ-4279 is similar to ML323 (Figure S2). The 4-cyclopropyl-6-methoxypyrimidin-5-yl group of KSQ-4279 occupies the same position as the 2-propan-2-ylphenyl group in ML323. The 1-propan-2-yl-4-(trifluoromethyl)imidazol-2-yl moiety is slightly shifted compared to the triazol-1-yl group of ML323 (RMSD of ring atoms ∼0.9 Å). The phenyl group of KSQ-4279 is rotated with respect to the ML323 structure, however, given the limits of the resolution, this is within the uncertainty of the modelling. N160 of USP1 is within hydrogen bonding distance of KSQ-4279 (Figure 1D), similar to the ML323 structure^29^. The β-turn on which the catalytic aspartates, D751 and D752^34^, reside, is pushed by the pyrazolo[3,4-d]pyrimidine group of KSQ-4279, slightly displacing these residues with respect to the inhibitor-free USP1 (Figure 2A). In ML323, the methyl substituent of the pyrimidine also pushes this β-turn but to a greater extent – also displacing S753. F101 is pushed out slightly to accommodate the methoxyl substituent of KSQ-4279 which was not observed for ML323-bound USP1 (Figure 2B). Overall, there are subtle changes in the cryptic binding site between KSQ-4279 and ML323.

**Figure 2:**
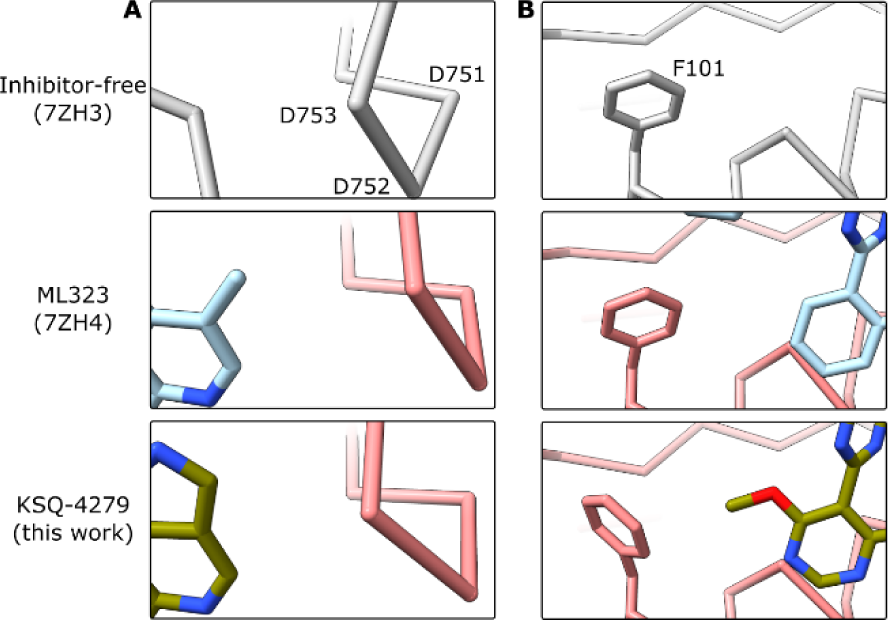
Differences in the effects of ML323 and KSQ-4279 on USP1 structure. (A) KSQ-4279 perturbs the β-turn, on which the catalytic aspartates D751 and D752 reside, to a lesser extent than ML323. (B) KSQ-4279 perturbs F101 whereas ML323 does not. Protein Cα atoms are shown.

The region with high structural heterogeneity in the ML323-bound structure^29^, spanning residues 168-195, also exhibits reduced order in the KSQ-4279-bound structure (Figure 3A). However, unlike the ML323 structure, helix α4 in this region remains in the same position as in the inhibitor-free structure, and the region between α3 and α4 is poorly resolved and was left unmodelled. We refer to residues 168-195 as the Mobilized by Inhibitor Region (MIR). Comparing the KSQ-4279 and ML323 maps, the density in the ML323 map resembles a combination of that observed in the KSQ-4279 structure and a conformation where α4 has slipped (Figure S3). We therefore sought to identify subsets of particles in our previous ML323 dataset (EMPIAR-11299^29^) in which the MIR was sufficiently resolved. We managed to isolate a subset of particles from which we could build an atomic model (ML323^subset^; Figure S4). In the ML323^subset^ structure, helix α4 slips closer to the inhibitor binding site by approximately 1.5 turns and the region between helix α3 and α4 (including β2) reorders to form another helix (Figure 3B). In both inhibitor-bound structures, residues near G194 are structurally heterogenous. A similar region in USP7 also exhibits plasticity and has been referred to as the switching loop^35^. This short stretch (residues 192-195 in USP1) may be innately flexible in the USP fold.

**Figure 3:**
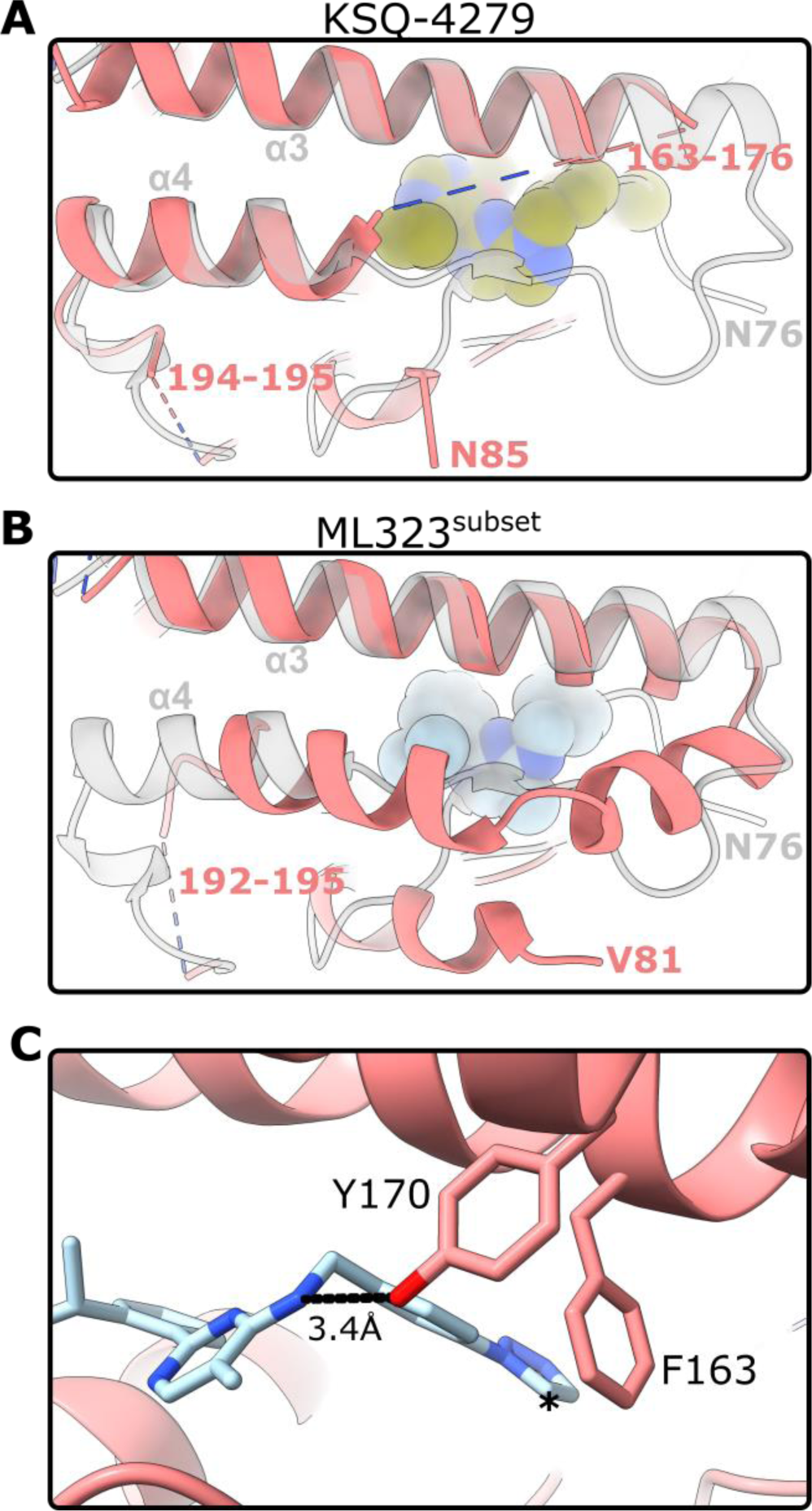
Comparison of the USP1 MIR between ML323 and KSQ-4279. (A) KSQ-4279 disrupts β-strands of the inhibitor-free state leaving residues 163-176 disordered. The inhibitor-free structure (7ZH3^29^) is shown in transparent grey. KSQ-4279 is shown as space-filling spheres based on van der Waals radii. (B) A subset of ML323-bound particles have reordered residues 167-191 resulting in formation of a new helix and slipping of helix α4, in addition to bending of helix α3. The inhibitor-free structure (7ZH3^29^) is shown in transparent grey. ML323 is shown as space-filling spheres based on van der Waals radii. (C) Additional interactions of the ML323^subset^ structure. Dashed lines indicate a potential hydrogen bond. Asterisk indicates where the isopropyl group of KSQ-4279 would clash with F163. ML323 is shown as sticks.

In the ML323^subset^ MIR, Y170 of the newly formed helix likely hydrogen bonds with the secondary amine joining the pyrimidine and benzyl groups, while F163 is brought close to the triazole group (Figure 3C). However, KSQ-4279 lacks a secondary amine in the equivalent position and has substituents that would clash with F163. As such, the conformation of the ML323^subset^ USP1 structure is incompatible with the KSQ-4279 molecule. This may explain the lack of order in this region when KSQ-4279 is bound. Aside from the major conformational changes required to establish the cryptic site, it appears KSQ-4279 binding maintains the USP1 fold in a slightly more native conformation than the binding of ML323.

In the absence of inhibitor there is a solvent accessible hydrophobic tunnel near the cryptic site (Figure 4A). Binding of ML323 or KSQ-4279 almost completely fills this tunnel. The tunnel is primarily plugged by the 2-propan-2-ylphenyl group in ML323 and the 4-cyclopropyl-6-methoxypyrimidin-5-yl group in KSQ-4279. Both inhibitors leave a small hydrophobic cavity, raising the possibility for expansion of the inhibitors into this cavity to improve binding affinity. The presence of this tunnel may facilitate preliminary binding to USP1 and direct conformational changes required to generate the cryptic site.

**Figure 4:**
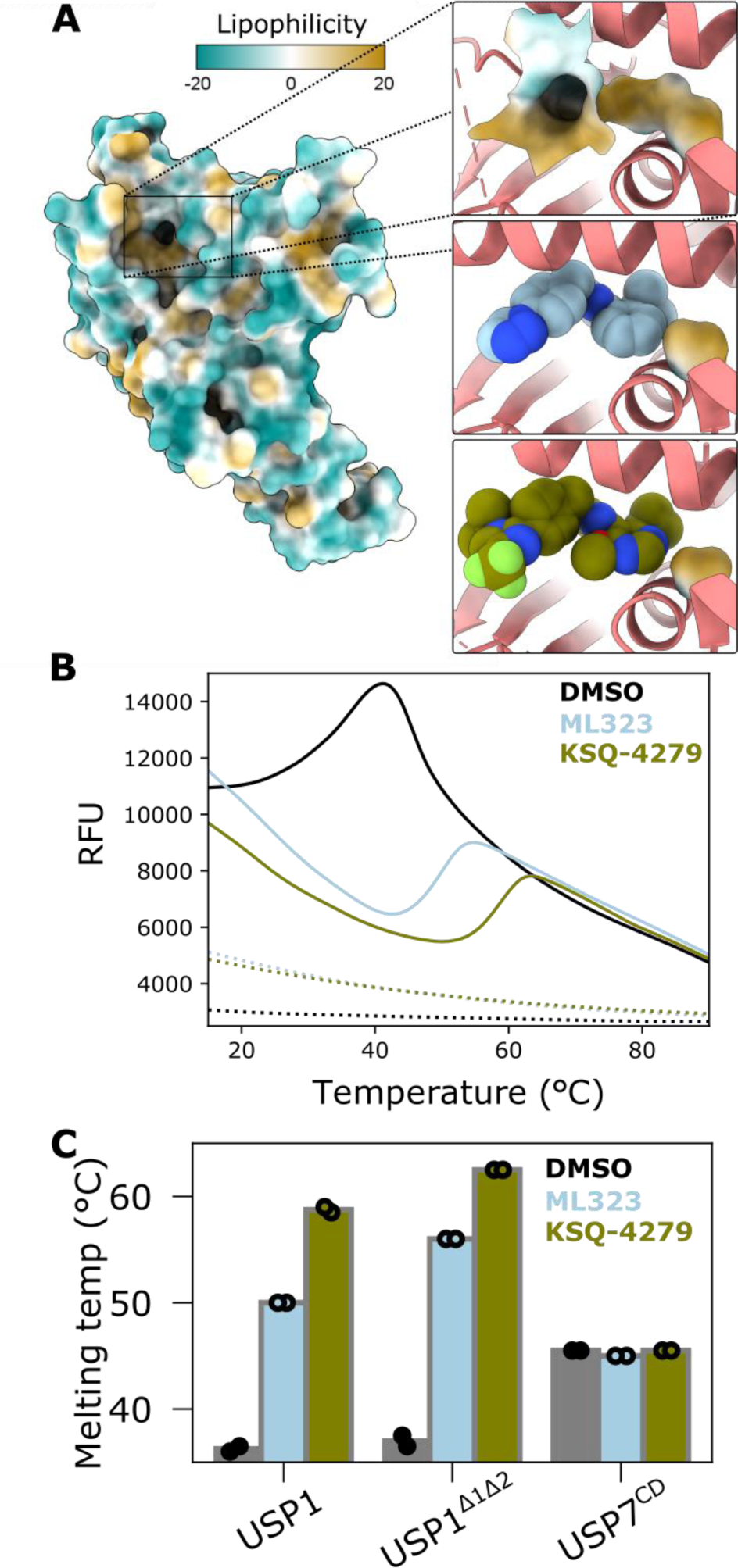
A hydrophobic tunnel is occupied by USP1 inhibitors. (A) Solvent accessible surface of inhibitor-free USP1 (7ZH3^29^) near to the cryptic binding site (top). KSQ-4279 and ML323 binding leaves only a small pocket remaining (middle and bottom, respectively). Solvent accessible pockets are shown as surfaces coloured by lipophilicity and inhibitors as space-filling spheres based on van der Waals radii. Superposition of each inhibitor-bound structures onto the inhibitor-free structure (7ZH3^29^) was performed. (B) Thermal shift assays of USP1 (3.8 μM) in the presence or absence of ML323 or KSQ-4279 (50 μM) (solid lines). ML323, KSQ-4279, and DMSO without USP1 are shown as dashed lines. (C) Quantification of thermal shift assays using the inflection point to estimate the melting temperature. USP7 catalytic domain (USP7^CD^) was included as a negative control. Two technical replicates for each USP were performed and are shown as circles.

Using thermal shift assays, we compared the effect of inhibitors on the isolated USP1 protein (Figure 4B-C). Strikingly, an excess of ML323 and KSQ-4279 increased the melting temperature by 14°C and 23°C, respectively. Both full-length USP1, and USP1 with insert 1 and 2 deleted (USP1^Δ1Δ2^) were shifted by similar extents (Figure 4C). This is consistent with stabilization of the USP fold itself rather than perturbing interaction with inserts 1 and 2. In contrast, the USP7 catalytic domain, used here as a control, did not show any change in melting temperature in the presence of either inhibitor. Overall, this is consistent with the inhibitors filling the hydrophobic tunnel to create a more stable hydrophobic core of USP1.

### Structural implications on selectivity of the inhibitors

Given the improved selectivity of KSQ-4279 over ML323 for USP12 and USP46 (Figure 1A), we sought to rationalise this from the structures. We aligned USP12^33^ and USP46^31^ to inhibitor-free USP1 and superposed the KSQ-4279 and ML323-bound structures onto these (Figure S5). We looked for protein-inhibitor clashes in the regions that remain fully ordered in USP1 upon inhibitor binding, i.e. excluding the MIR and RIR. ML323 clashes with N97 and the β-turn on which the catalytic aspartates reside^34^ in all three USPs. KSQ-4279 clashes with N97 in all three USPs and the same β-turn as ML323 clashes in USP12. A further clash for KSQ-4279 is observed in all superpositions between the methoxyl group and F101 (F59 and F55 in USP12 and USP46, respectively). We therefore propose that rearrangement of F101 in USP1 (Figure 2B), cannot be as easily accommodated in USP12 and USP46, as a mechanism for the increased selectivity of KSQ-4279 over ML323. However, this mechanism assumes the region homologous to the USP1 MIR also becomes flexible in USP12 and USP46. We cannot rule out that interactions between the 1-propan-2-yl-4-(trifluoromethyl)imidazol-2-yl group and the regions homologous to the MIR, which are much shorter in USP12 and USP46, mediate the difference in selectivity between ML323 and KSQ-4279.

We further sought to identify how USP1, USP12, and USP46 may be selected for over other USPs. We hypothesised that the tunnel accommodating the 4-cyclopropyl-6-methoxypyrimidin-5-yl group of KSQ-4279 may be present to some extent in USP12 and USP46 but not other USPs. We aligned 48 alphafold structures and the KSQ-4279 structure by the USP fold and looked for cavities near the 4-cyclopropyl-6-methoxypyrimidin-5-yl group (Figure S6). Although USP12 and USP46 do have small cavities in this region, so too does USP11 which was not inhibited by either compound (Figure 1A), therefore presence of a hydrophobic tunnel in this region does not appear to exclusively mediate the selectivity.

### Effect of inhibitors on catalysis

The catalytic site of the KSQ-4279-bound structure was perturbed as previously observed for ML323^29^ (Figure 5A). The β-turn on which D751 and D752 reside, is displaced; however, the density for the carboxyl groups of D751 and D752 is not sufficient to unambiguously assign these atoms in the KSQ-4279-bound structure. Regardless, the hydrogen bonding between H593 and D751 is disrupted. Recently, both D751 and D752 have been demonstrated to be important for USP1 catalysis^34^. Therefore, inhibition by KSQ-4279 and ML323 may not be as simple as our previous hypothesis of stabilizing a flipped H593 conformation that cannot efficiently deprotonate C90, reducing the catalytic cysteines ability to exert a nucleophilic attack at the isopeptide bond^29^. Indeed, N85 is involved in stabilization of the oxyanion intermediate during catalysis for other USPs^36,37^ and it forms a hydrogen bond with D752 in USP1 in the absence of any inhibitor (Figure 5A). Binding of either inhibitor not only disrupts this bonding but displaces N85 altogether. As such, inhibitor binding may not only impede the coordination of H593 but also destabilize the oxyanion intermediate.

**Figure 5:**
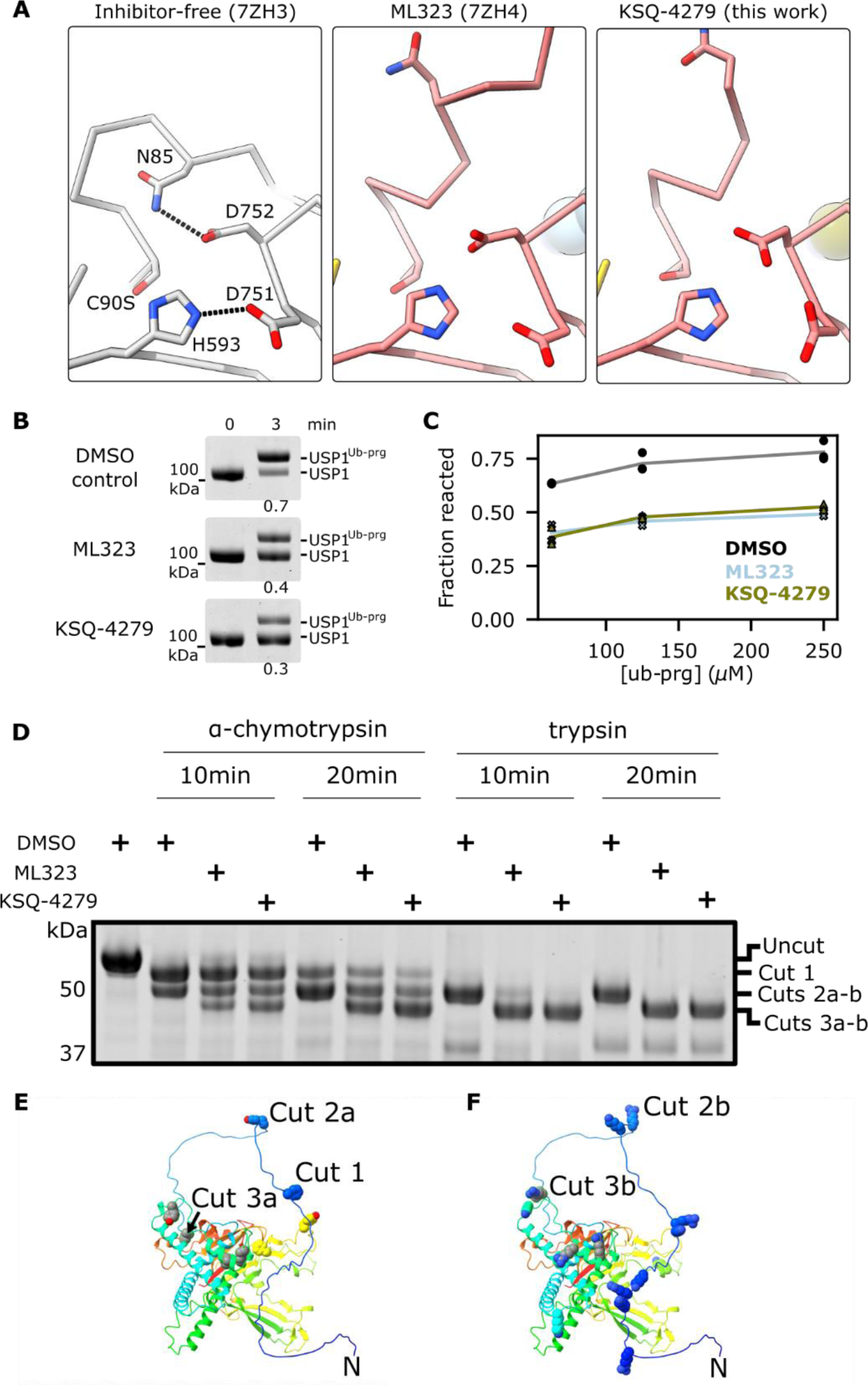
Disruption of the USP1 active site. (A) ML323 or KSQ-4279 binding disrupts hydrogen bonding of the catalytic aspartates and displaces the oxyanion stabilizing residue, N85. Hydrogen bonds of the inhibitor-free structure are shown as dashed lines. (B) Gel-based assay for reactions of USP1 (2 µM) in the presence or absence of ML323 or KSQ-4279 (25 µM) with Ub-Prg (6 μM) at room temperature. Fraction of the total signal corresponding to the reacted band is given below each reaction. Two technical replicates were performed. (C) Gel-based assay of reactions of 2 µM USP1 alone or with 25 µM inhibitor with excess Ub-Prg on ice using a single time-point. Fraction reacted was quantified using densitometric analysis of the bands. At least two technical replicates were performed. (D) Limited proteolysis reactions of USP1^Δ1Δ2^ with different proteases. (E) AlphaFold model of USP1^Δ1Δ2^ with residues of highly probable α-chymotrypsin cleavage sites and high solvent accessibility shown as spheres. Residues with high cleavage probability that are only accessible in inhibitor-bound structures are colored grey. The N-terminus and probable sites for cleavage products generated by cuts 1-3 are indicated. (F) AlphaFold model of USP1^Δ1Δ2^ with residues of highly probably trypsin cleavage sites and high solvent accessibility shown as spheres. Protein cartoon is colored by Jones’ rainbow. Residues with high cleavage probability that are only accessible in inhibitor-bound structures are colored grey. The N-terminus and probable sites for cleavage products generated by cuts 2 and 3 are indicated.

To examine the effect of inhibitor on the early stages of catalysis we used ubiquitin-propargyl amide (Ub-Prg) reactions. The catalytic cysteine of deubiquitinases can nucleophilically attack the Ub-Prg probe to yield a stable covalent adduct^38,39^. Reaction of Ub-Prg with inhibitor-free USP1 at room temperature was ∼70% complete within 3 minutes (Figure 5B). In the presence of an excess of both inhibitors, the reaction between Ub-Prg and USP1 was reduced, with ∼30-40% USP1 reacted at 3 minutes. The reduced reaction with Ub-Prg in the presence of inhibitor at 3 minutes may be due to reduced Ub-Prg binding rate or reduced reaction rate. To increase the Ub-Prg binding rate such that this step is not rate-limiting we used >50 μM of Ub-Prg (>30-fold molar excess over USP1). We performed reactions on ice to enable single time-point measurements at 3 minutes. Both ML323 and KSQ-4279 reduced the reaction with Ub-Prg to a similar extent under these conditions (Figure 5C). This result is consistent with binding of ML323 or KSQ-4279 perturbing nucleophilic attack of the catalytic cysteine, however, the oxyanion hole may also be important in this reaction^40^.

The oxyanion stabilizing residue, N85, is located on the RIR of USP1. We hypothesize that its displacement upon binding of ML323 or KSQ-4279 would make the RIR susceptible to more generic proteases. Therefore, we explored limited proteolysis assays to characterize inhibitor binding (Figure 5D, Figure S7). We treated USP1^Δ1Δ2^ with α-chymotrypsin or trypsin and found that the major product was truncated to two slightly smaller products in the absence of inhibitor. In the presence of ML323 or KSQ-4279 an additional, fast migrating band was also present, indicating further truncation. Considering the cleavage sites preferences for α-chymotrypsin and trypsin, and the accessibility of the sites based on an AlphaFold model of USP1^Δ1Δ2^ (Figure 5E-F, Figure S8), we interpret the second species to be cleavage at Y52 (Cut 2a) or R53 (Cut 2b) for α-chymotrypsin and trypsin, respectively, and the third species to be cleavage at F80 (Cut 3b) or R74 (Cut 3a) for α-chymotrypsin and trypsin, respectively. Cuts 3a-b only occur in the presence of ML323 or KSQ-4279 which is consistent with inhibitor binding displacing the RIR containing N85. We do not observe significant cleavage at the R74, F80 sites in the absence of inhibitor, suggesting this region is not significantly exposed in the absence of inhibitor and consistent with an induced fit mode of binding within the hydrophobic tunnel.

### Limitations

One of the limitations of this study is that the structures of inhibitor-bound USP1 were determined in the context of the C90S mutation of USP1, and USP1 is bound to substrate, which may perturb the enzyme structure compared to substrate-free wild-type enzyme. However, we were unable to obtain reliable structures of USP1-UAF1 alone with the inhibitor. Despite the presence of substrate, the extensive inhibitor interactions at the cryptic binding site are almost certainly similar in the absence of substrate. As our data contained a mixture of inhibitor-bound and inhibitor-free states we used classification algorithms to sort the particles. However, cryo-EM data is inherently noisy and these algorithms are imperfect, therefore there is potential for misclassification to perturb the resulting reconstructions. We explored many classification parameters to mitigate this. In the KSQ-4279 structure, the density for N97, adjacent to the cryptic site is poor. This may be a manifestation of misclassified particles, additional sub-states or possible radiation damage. Finally, the reconstruction resolutions of the ML323 and KSQ4279 structures vary, which complicates comparison of the maps. In particular we are unable to reliably model waters in the KSQ-4279 structure. For comparison of the MIR, the maps were low-pass filtered to 5 Å to mitigate the effect of the difference in resolutions.

## Conclusions

Overall, our data suggest that the clinical USP1 inhibitor, KSQ-4279, and the well-established tool compound ML323 exert similar inhibitory effects on USP1. Both inhibitors bind to a cryptic binding site in a similar mode (Figure 6). However, the surrounding residues of the protein are disrupted in different ways. KSQ-4279 is clearly more selective than ML323 with respect to its impact on other USP enzymes. The observation that USP12 and USP46 are inhibited at relatively high concentrations of the ML323 inhibitor is consistent with selectivity data for another USP1 inhibitor, I-138^24^. KSQ-4279 may exhibit greater selectivity for USP1 over USP12 and USP46 compared to ML323 due to the methoxyl substituent of KSQ-4279 further disturbing the binding site, however this requires further investigation. The occupancy of a hydrophobic tunnel by the inhibitors appears to drive a substantial increase in thermal stability of USP1. This work helps to define the mechanism of action of the USP1 inhibitor KSQ-4279, currently in clinical trials.

**Figure 6:**
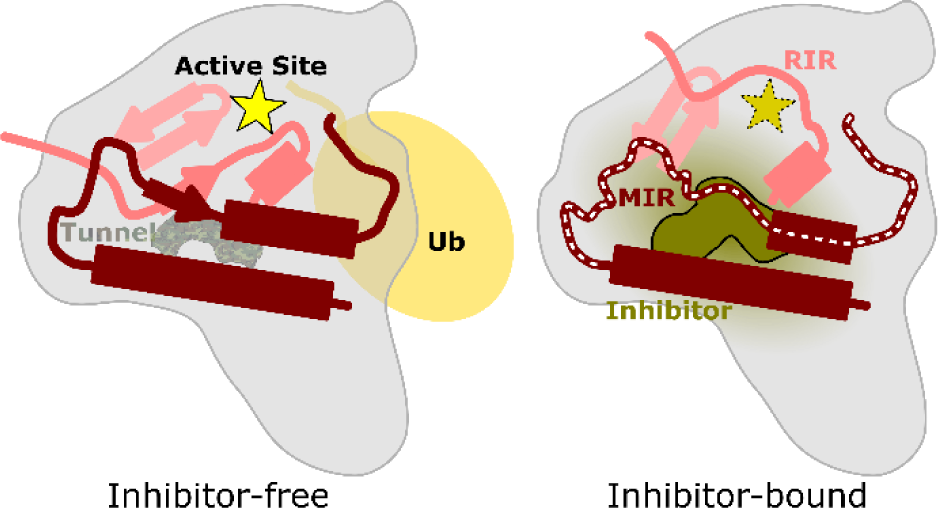
Schematic representation of the effect of inhibitor binding on USP1 structure. In the inhibitor-free state, USP1 has a hydrophobic tunnel and ordered active site, while inhibitor binding disrupts the active site, MIR (white dashes), and RIR.

## Acknowledgements

We thank Kevin Parkes and Rishi R. Shah from Ubiquigent for comments on the manuscript. We thank past and current members of the Walden laboratory for experimental suggestions, comments on the manuscript and their support. We acknowledge the Scottish Centre for Macromolecular Imaging (SCMI) for access to cryo-EM instrumentation, funded by the MRC (MC_PC_17135, MC_UU_00034/7) and SFC (H17007). Cryo-EM data were collected at eBIC via the Industry Access route. We thank Dr. Rachel Toth for expression plasmids. We thank Mark Meenan, Paul McLaughlin, and Iain Sim for maintenance of the GPU server running cryoSPARC.

## Author contributions (CRediT)

Conceptualization – M.R., M.G., S.F., H.W.; Investigation – M.R., M.G., C.A., S.L.; Supervision – S.F., H.W.; Data Curation – M.R.; Visualization – M.R.; Writing (original draft) – M.R.; Writing (review and editing) – M.R., M.G., C.A., S.F., S.L., H.W.

## Data and materials availability

All data needed to evaluate the conclusions in the paper are present in the paper and/or the Supplementary Materials. The atomic coordinates have been deposited to the PDB under accession codes 9FCI and 9FCJ. The cryo-EM maps have been deposited to the EMDB under accession codes EMD-50316 and EMD-50317.

## Methods

### Protein expression and purification

Proteins, all human homologs, were expressed in either insect cells or bacteria as described previously^3,41^. Protein purification buffers and columns used are provided in Table S2. Briefly, His_6_-TEV-USP1^G670A,G671A^, His_6_-TEV-USP1^G670A,G671A,C90S^, His_6_-TEV-USP1^Δ229-408,Δ608-737^, His_6_-3C-UAF1, His_6_-3C-FANCD2, His_6_-TEV-V5-FANCI, and His_6_-TEV-V5-FANCI^S556A,S559A,S565A^ were expressed separately in *Sf*21 insect cells. Cells were lysed by sonication, clarified, and purified by Ni-NTA affinity then anion exchange chromatography. At this stage protein aliquots were occasionally flash frozen in liquid nitrogen and stored -80°C. For His_6_-TEV-USP1^G670A,G671A^, His_6_-TEV-USP1^G670A,G671A,C90S^ and His_6_-TEV-USP1^Δ229-408,Δ608-737^, TEV protease treatment was performed overnight at ∼1:10 protease to target protein with gentle agitation, resulting in cleaved protein with an N-terminal glycine extension. Subtractive Ni-NTA affinity chromatography was subsequently performed. All proteins were concentrated to 5-10 mg/mL and separated by gel filtration. Purified protein was concentrated to 5-15 mg/mL, flash frozen, and stored at -80°C in 10-20 μL single-use aliquots. All steps were performed on ice or at 4°C and completed within 24-36 hours of lysis. FANCD2 was ubiquitinated and purified using an engineered Ube2T and SpyCatcher-SpyTag setup described in detail elsewhere^42^. For FANCD2, the His_6_-3C tag was removed by 3C protease treatment during preparation of the mono-ubiquitinated version.

The USP domain (residues 208-560) from USP7 was expressed as a His_6_-smt3 fusion in BL21 *E.coli* cells. Cells were grown to an OD600 of ∼0.6 and expression induced with 0.1 mM IPTG at 16°C for ∼18 hours. Cells were harvested and lysed by sonication. Lysate was clarified and bound to Ni-NTA resin. His_6_-ULP1 protease was added to the resin and incubated overnight at 4°C to cleave the His_6_-smt3 tag. Flow-through was collected and purified by anion exchange chromatography, followed by gel filtration. Purified protein was concentrated to ∼10 mg/mL, flash frozen, and stored at -80°C in 10-20 μL single-use aliquots.

Ubiquitin-propargylamine (Ub-Prg) was prepared with an N-terminal twin-strep tag. Ubiquitin^1–75^-Intein-Chitin binding domain (CBD) plasmid was gifted from Dr Yogesh Kulathu (University of Dundee) and a twin-Strep and a 3C cleavage site was cloned N-terminally. Twin-strep-3C-ubiquitin^1–75^-Intein-CBD was produced in BL21 *E.coli* cells. Cells were grown to an OD600 of 0.4-0.5 and expression induced with 0.3 mM IPTG at 16°C for ∼24 hours. Cells from 6 L expression culture were harvested, lysed by sonication, clarified, and bound to chitin beads. Beads were washed with ∼40x bead volume of Wash Buffer 1 followed by 10x bead volume of Wash Buffer 2. Intein auto-cleavage was performed in three elution steps, each with >2.5x bead volume of Elution Buffer and >24 hours incubation. Eluant was concentrated to ∼20 mg/mL and pH adjusted to 8.0 with 0.1 M NaOH (or dialysed into 50 mM HEPES). The propargylamine (prg) warhead was then conjugated by incubating with 0.25 M propargylamine 4-6 h at 16°C in the dark and mild agitation at 30 min intervals. Ub-Prg was concentrated and separated by SEC using an SD75 16/600 column in 1x PBS before concentrating to ∼10 mg/ml and storage at -80 °C.

Protein concentrations were determined using the predicted extinction coefficients at 280 nm^43^ and absorbance via a Nanodrop. The ratio of 260nm/280nm was ≤0.65 for all protein batches used in subsequent experiments.

### Ubiquitin-rhodamine assays

To determine the potency of both compounds on USP1, the ability of USP1-UAF1 (0.008 nM) to cleave the ubiquitin-rhodamine substrate (100 nM) was determined in the presence or absence of a half-log, 8-point dilution series of each compound. IC_50_ values were determined from the dose response curve of each compound by fitting the Hill equation.

The selectivity of KSQ-4279 and ML323 was then evaluated in the DUB*profiler*™ assay (Ubiquigent) against 48 individual deubiquitinase enzymes, at concentrations of 0.01, 0.1 and 1, 10 and 100 µM of each compound, the latter being equivalent to at least 10,000-fold greater concentrations than the IC_50_ against the primary target.

Values in the presence of compound were compared with DMSO controls to give the percentage of remaining activity. Neither compound showed any significant autofluorescent properties in the ubiquitin-rhodamine assay up to 100 µM.

### Cryo-EM sample preparation

The USP1^C90S^-UAF1-FANCI-FANCD2^Ub^ complex was prepared by mixing the four individually purified subunits at 5:5:1:1 (USP1^G670A,G671A,C90S^:His_6_-3C-UAF1:His_6_-TEV-V5-FANCI^S556A,S559A,S565A^:FANCD2^Ub^) as described previously^29^. The complex was exchanged into EM buffer (20 mM Tris pH 8.0, 150 mM NaCl, 2 mM DTT) using a Bio-Spin P-30 column (Bio-Rad). The concentration of complex was estimated from absorbance at 280 nm (assuming no loss of any of the protein components). 1.2 equivalents of dsDNA per FANCI-FANCD2^Ub^ was then added (61 base pairs; TGATCAGAGGTCATTTGAATTCATGGCTTCGAGCTTCATGTAGAGTCGACGGTGCTGGGAT; IDT). Finally, KSQ-4279 was added at 2 equivalents of USP1-UAF1. The sample was incubated at room temperature for 5 minutes immediately prior to preparing grids. UltrAuFoil R1.2/1.3 300 mesh grids were glow discharged at 35 mA for 60 secs. A 3.0 μL aliquot of 9.6 μM USP1-UAF1, 1.9 μM FANCI-FANCD2^Ub^, 2.3 μM dsDNA, 18.8 μM ML323 was applied. The grids were blotted for 3.0 secs and vitrified in liquid ethane using a Vitrobot (ThermoFisher) operating at ∼95% humidity at 15°C.

### Cryo-EM sample data collection and processing

Grid screening and data collection was performed on a Titan Krios (ThermoFisher) located at eBIC (Diamond Light Source) equipped with a K3 detector and BioQuantum Energy Filter (Gatan). A total of 6,581 movies were collected using Beam-Image shift. Movies were collected in Counted Super Resolution with 2x binning and CDS at a pixel size of 0.83 Å using EPU (ThermoFisher). An energy filter slit width of 20eV was used. Movies were collected with a total dose of ∼62 e^-^/Å^2^ over 50 frames at a rate of ∼6 e^-^/px/sec.

Subsequent processing was performed in cryoSPARC v3.3 and v4.4^44^ (Figure S1). Patch motion correction, patch CTF estimation, and manual curation was performed resulting in 5,675 dose-weighted, motion corrected micrographs. Both template picking and topaz^45^ were used to identify potential particles which were extracted at 384×384 pixels Fourier cropped to 128×128. Multiple 2D classifications and heterogenous refinement were used in parallel for initial cleaning with selected classes pool and duplicates removed. Further particle cleaning was performed using heterogenous refinement with one good starting model and 5 “junk” starting models distinct from the protein complex of interest, all low pass filtered to 20 Å. A final round of heterogenous refinement was performed using the same starting model low-pass filtered at 12 Å, and two copies at 30 Å. Five full passes through the particle sets, after O-EM iterations, were used in heterogenous refinements. Particles corresponding to the highest resolution class subjected to another round of duplicates removal and were then re-extracted with a boxsize of 384×384 pixels without Fourier cropping. Non-uniform refinement^46^ yielded a structure with a global resolution of 3.6 Å from 135,680 particles. These were subjected to reference-based motion correction and further non-uniform refinement yielding a structure with a global resolution of 3.5 Å from 135,646 particles. Local refinement was performed with a mask covering USP1 and ubiquitin using a gaussian priors of 3° over rotation and 2 Å over shifts with marginalization and non-uniform refinement. 3D classification was performed with a smaller mask covering the cryptic binding site and adjacent regions. Classes with density consistent with inhibitor binding were pooled and subjected to another round of local refinement with a mask covering USP1 and ubiquitin.

For reprocessing of the previously published dataset with ML323 (EMPIAR-11299)^29^, reference-based motion correction was performed on particles from the consensus reconstruction prior to local motion correction (Figure S4). Local refinement was performed with a mask covering USP1 and ubiquitin using a gaussian priors of 3° over rotation and 2 Å over shifts with marginalization and non-uniform refinement. 3D classification was performed with a mask covering the inhibitor binding site and 10 classes using “simple” initialization. A class with interpretable density for residues 168-191 (ML323^subset^) was passed through local refinement again with the mask covering USP1 and ubiquitin.

### Model building and refinement

KSQ-4279 and ML323^subset^ structures were built using USP1 and ubiquitin using the previous structure of ML323-bound structure (PDB ID: 7ZH4) as an initial model. Manual model editing was performed using COOT^47^ and ISOLDE^48^. KSQ-4279 restraints were calculated using the GRADE web server (http://grade.globalphasing.org/). Automated refinement against the globally sharpened maps, with B-factors estimated from the Guinier plot, were performed using phenix real-space refinement^49^. A refinement resolution of 3.8 Å and 3.2 Å was used for the KSQ-4279 and ML323^subset^ structures, respectively. Bond and angle restraints for the USP1 Zinc finger, as well as Ramachandran restraints, were incorporated during automated refinement. Cryo-EM data and model statistics are reported in Table S1. The FSC between the models and maps were computed using phenix^50^ (Figure S1, S4). Structures and maps were aligned, analyzed, and figures produced, using ChimeraX^51^.

### Deubiquitination assays

Deubiquitination reactions were performed as described previously^29^. A 2x substrate mix and a 2x enzyme mix were prepared separately and mixed 1:1 to initiate the reaction. Both mixes were setup on ice, and then incubated at room temperature for at least 20 min prior to reaction initiation and during the reaction.

The 2x substrate mix was prepared by diluting stocks of FANCD2_Ub_ (>29 μM), His_6_-V5-TEV-FANCI (>50 μM), and dsDNA (100 μM; 61 base pairs) with DUB buffer (20 mM Tris pH 8.0, 75 mM NaCl, 5% glycerol, 1 mM DTT). The resulting 2x mix was composed 2 μM FANCD2_Ub_, 2 μM FANCI, 8 μM dsDNA. The 2x enzyme mixes were prepared by diluting concentrated stocks (≥30 μM) of USP1, His_6_-3C-UAF1, and ML323 or KSQ-4279 or DMSO control with DUB buffer. The resulting 2x enzyme mixes were composed of 200 nM USP1^G670A,G671A^, 200 nM UAF1, 1% DMSO, and 50 μM inhibitor, where included. Aliquots of 4 μL of reaction were terminated at the indicated timepoints by addition of 20 μL 1.2x NuPAGE LDS buffer (Thermo Fisher) supplemented with 120 mM DTT. SDS-PAGE was performed using Novex 4–12% Tris-glycine gels (Thermo Fisher) and subsequent staining of the gels with Instant-Blue Coomassie stain (Expedeon).

### Thermal shift assays

Concentrated stocks of USP1^G670A,G671A^, USP1^Δ229-408,Δ608-737^ (USP1^Δ1Δ2^), or USP7^208-560^ (USP7^CD^) (>100 μM) were diluted to 4 μM with TSA buffer (50 mM Tris pH 8.0, 200 mM NaCl, 5% glycerol, 1 mM DTT). This was added to SYPRO Orange and inhibitor or DMSO control to yield a final concentration of 3.8 μM protein, 5x SYPRO Orange, 50 μM inhibitor, 1% DMSO. Thermal shift assays were performed using a CFX96 Touch Real-Time PCR Detection System (Biorad) with 50 μL of sample heated from 15 °C to 90 °C at 3 °C/min and with fluorescence measured at 0.5 °C intervals. Data were analyzed using CFX Maestro (Biorad) and plotted with matplotlib^52^. Subtraction of controls without protein from the melting curves did not affect estimation of melting temperature.

### Ubiquitin-prg reactions

A 2x stock of USP1^G670A,G671A^ or USP1^Δ1Δ2^ was prepared containing 4 μM protein, 50 μM inhibitor, 1% DMSO in prg reaction buffer (1x PBS supplemented with 0.4mM TCEP). A 2x stock of Ub-Prg was prepared in prg reaction buffer, with the concentration of Ub-Prg varying depending on the final assay concentration. For reactions on ice, all components apart from inhibitor were prepared on ice. To initiate reaction, the two stocks were mixed 1:1. Reactions were terminated at indicated timepoints by mixing with an equal volume of 2x NuPAGE LDS buffer (Thermo Fisher) supplemented with 200 mM DTT. SDS-PAGE was performed using Novex 4–12% Tris-glycine gels (Thermo Fisher) or in-house 9% Tris acrylamide gels. Subsequent staining of the gels was performed with Instant-Blue Coomassie stain (Expedeon).

### Limited proteolysis

Reactions were performed with 10 μM USP1^Δ1Δ2^ and 25 μM inhibitor or DMSO control in 20mM Tris pH 8, 150mM NaCl, 5% glycerol, 0.4mM TCEP, 0.5% DMSO with 0.005 mg/mL α-chymotrypsin or trypsin. Reactions were terminated at indicated timepoints by mixing with an equal volume of 2x NuPAGE LDS buffer (Thermo Fisher) supplemented with 200 mM DTT. SDS-PAGE was performed using Novex 4–12% Bis-Tris gels (Thermo Fisher) and subsequent staining of the gels with Instant-Blue Coomassie stain (Expedeon).

PeptideCutter^53^ (https://web.expasy.org/peptide_cutter/) was used to predict sites with >70% and >90% cleavage probabilities for α-chymotrypsin or trypsin, respectively. An AlphaFold^54^ model of USP1^Δ1Δ2^ was generated using ColabFold^55^ with one recycle and no templates or relaxation. ChimeraX was used to measure the solvent accessible surface area of the backbone atoms for each predicted site, those computed to be >20 Å^2^ were considered as potential cleavage sites.

