## Supplementary Figures and Tables for "Structural and biochemical insights on the mechanism of action of the clinical USP1 inhibitor, KSQ-4279"

### Supplementary Materials

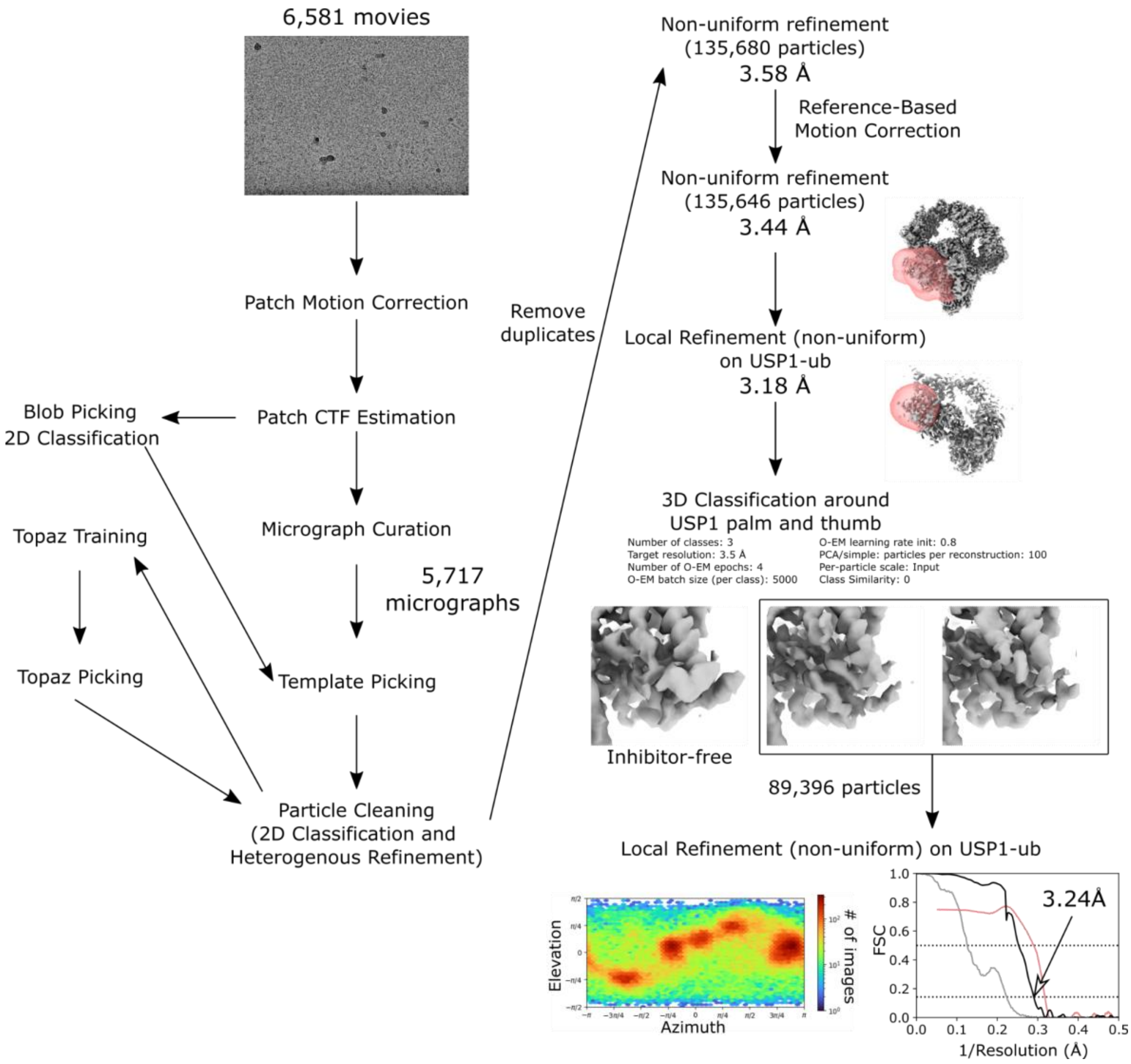


**Figure S1:** Cryo-EM particle processing workflow for the KSQ-4279-bound structure. Half-map FSCs for no mask (dark gray) and tight mask with correction by noise substitution (black) and distributions of particle orientations are shown for the final reconstruction.

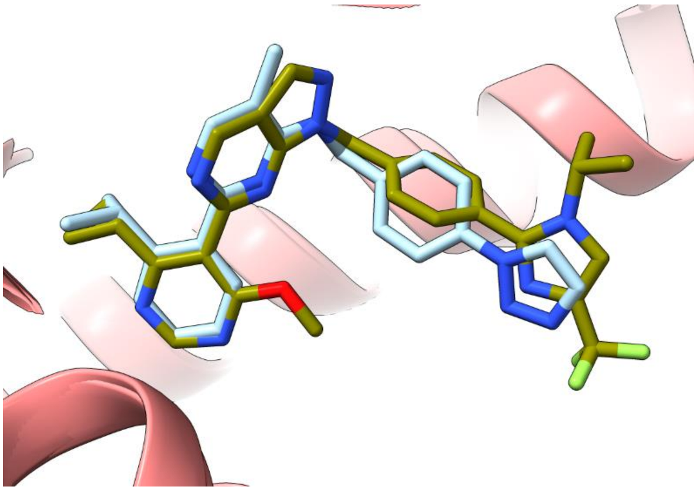

**Figure S2:** Superposition of the inhibitor binding sites for KSQ-4279 (olive) and ML323 (blue; 7ZH4).

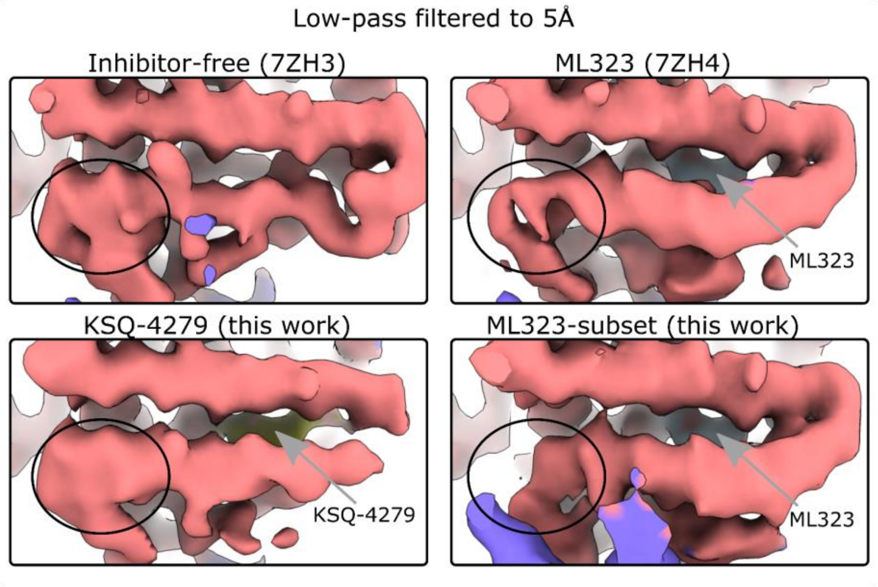


**Figure S3:** Comparison of the disordered region between ML323 and KSQ-4279 in the cryo-EM maps. Cryo-EM maps were low-pass filtered to 5 Å to mitigate differences in resolution. The circled region is similar between the inhibitor-free and KSQ-4279 structures, while with ML323 there are at least two conformations present.

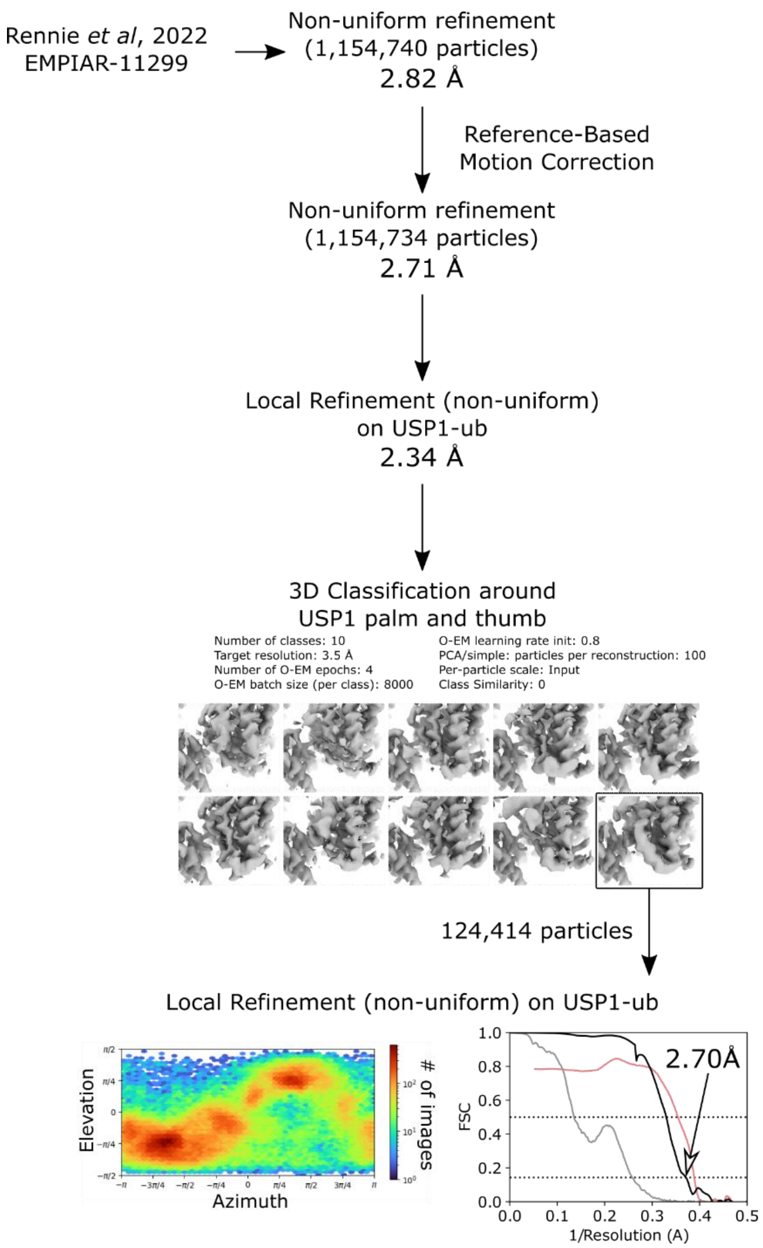


**Figure S4:** Cryo-EM particle processing workflow for the ML323$^{subset}$ structure. Half-map FSCs for no mask (dark gray) and tight mask with correction by noise substitution (black) and distributions of particle orientations are shown for the final reconstruction.

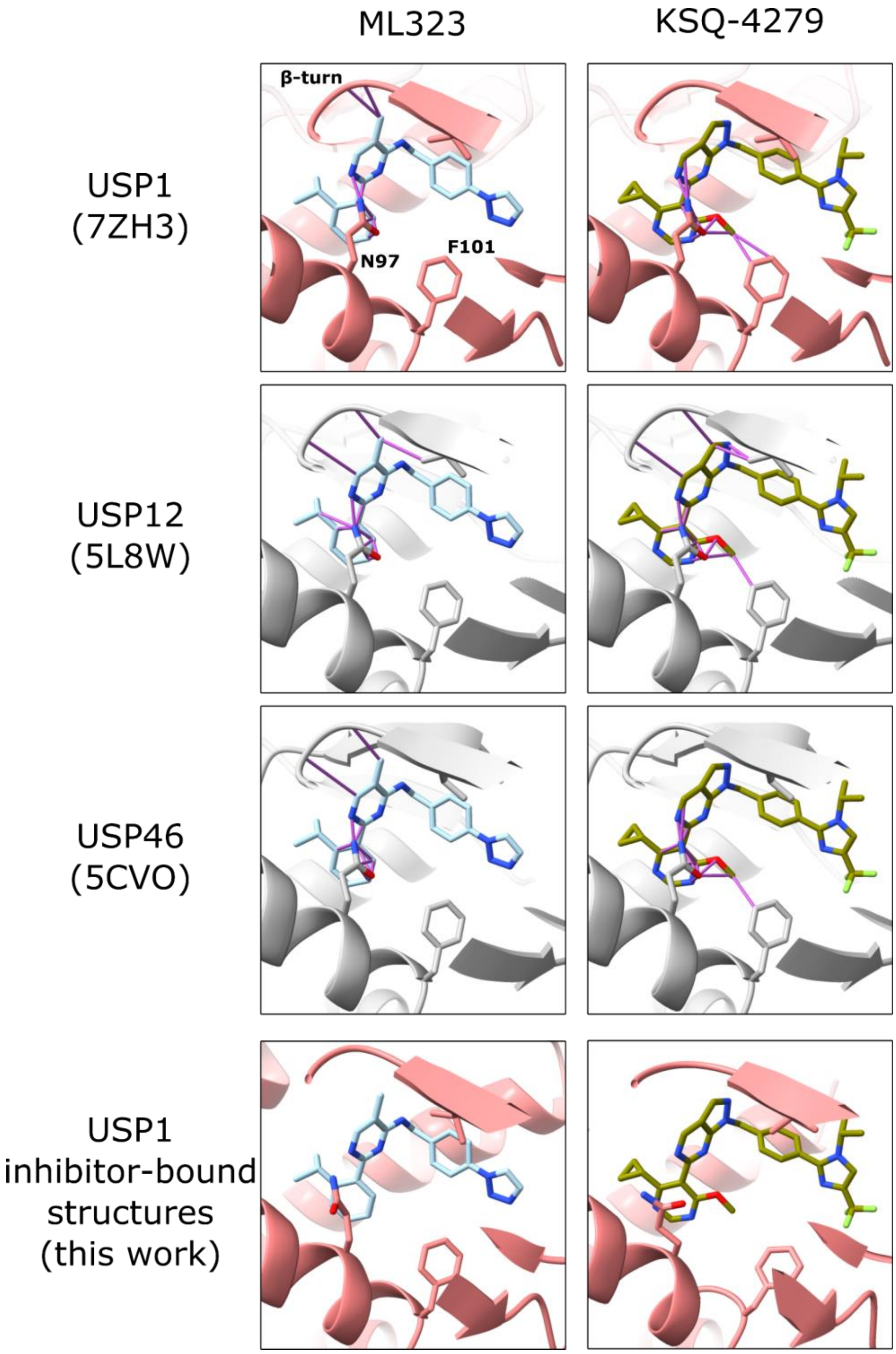


**Figure S5:** Superposition of the inhibitor-bound structures onto inhibitor-free USP1 (7ZH3[29]), USP12 (5L8W[33]) and USP46 (5CVO[31]) structures. The RIR and MIR are hidden. Clashes between non-RIR or -MIR residues are shown as purple lines.

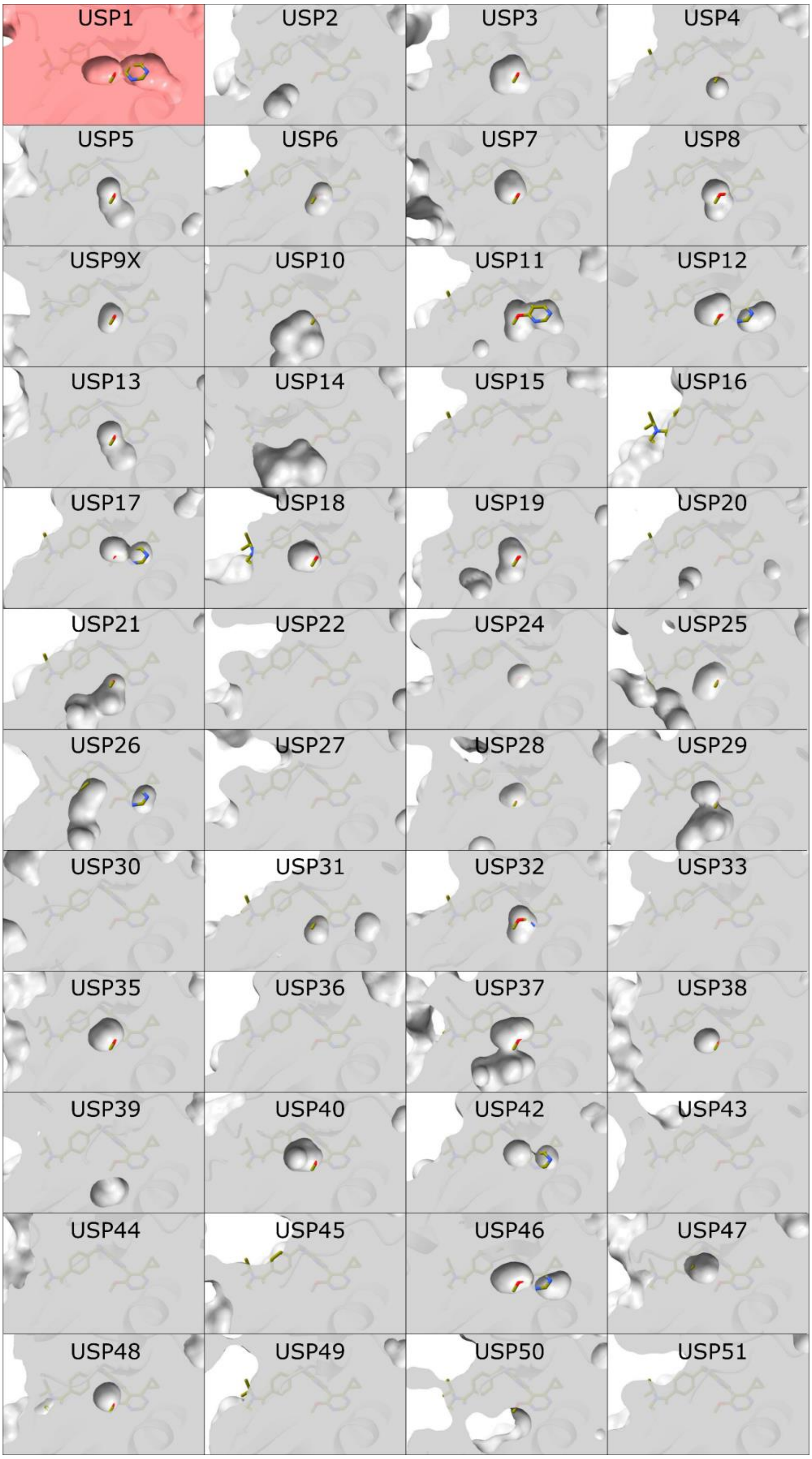


**Figure S6:** Superposition of the KSQ-4279-bound structure onto AlphaFold models of 48 USPs, focusing on the hydrophobic tunnel.

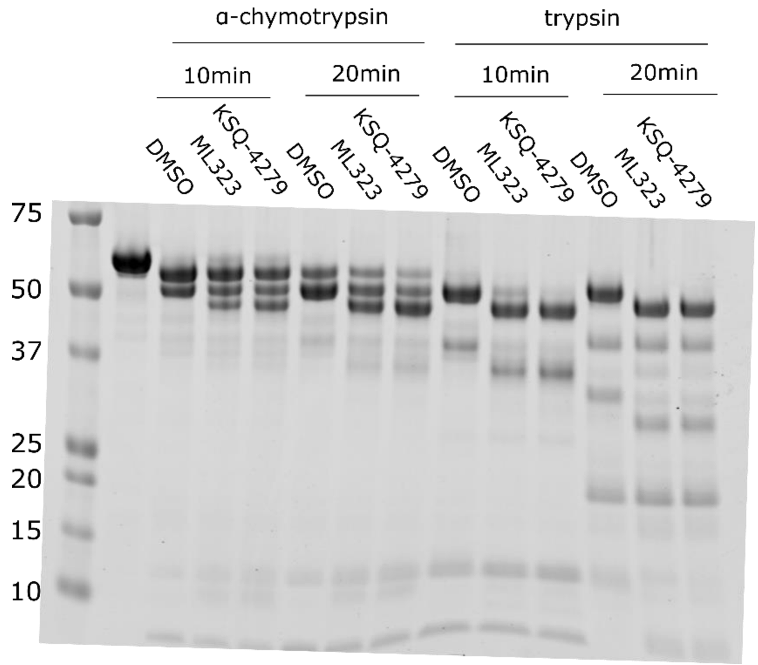


**Figure S7:** Limited proteolysis of USP1$^{\Delta1\Delta2}$.

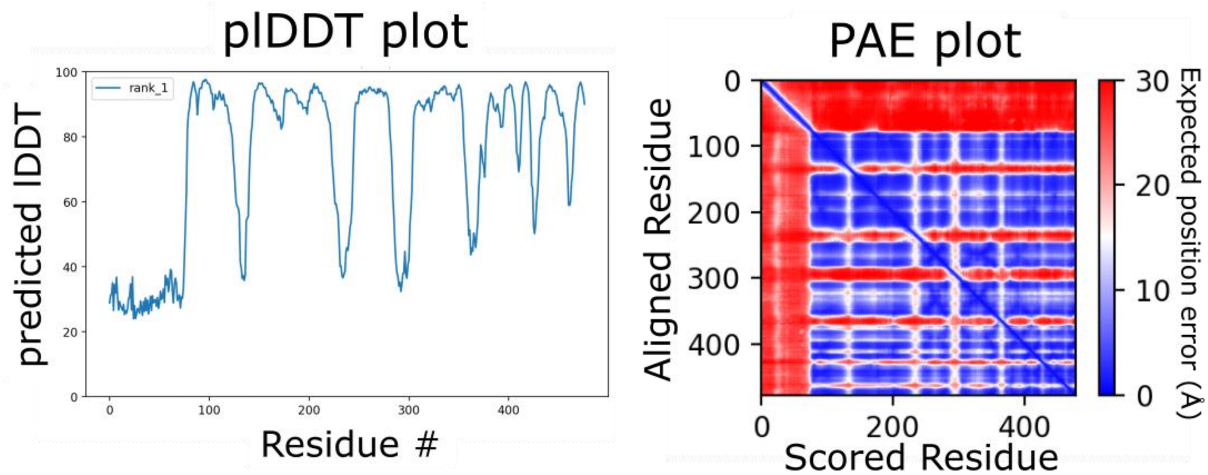


**Figure S8:** Confidence metrics for the USP1$^{\Delta1\Delta2}$ AlphaFold model.

**Table S1.** Cryo-EM data collection and model refinement statistics

| | KSQ-4279 Consensus (before classification) | KSQ-4279 Focused (after classification) | ML323[subset] Focused (EMPIAR-11299) |
|---|---|---|---|
| | | **Data collection and processing** | |
| Microscope | Krios | | - |
| Detector | K3 | | - |
| Nominal Magnification | 105,000x | | - |
| Voltage (kV) | 300 | | - |
| Electron Dose ($e^{-}/Å^2$) | ~60 | | - |
| Defocus range (μm) | ~0.5-2.0 | | - |
| Pixel Size (Å) | 0.83 | | 1.06 |
| Symmetry imposed | C1 | | C1 |
| Map resolution (Å) | 3.44 | 3.24 | 2.70 |
| FSC threshold | 0.143 | 0.143 | 0.143 |
| Map resolution range (Å)[a] | - | 3.1-4.0 | 2.5-3.5 |
| FSC threshold | - | 0.143 | 0.143 |
| EMDB ID | EMD-50316 (additional map) | EMD-50316 | EMD-50317 |
| | | **Refinement** | |
| Initial models used | - | 7ZH4 | 7ZH4 |
| Map sharpening B-factor ($Å^2$) | - | 97.9 | 81.5 |
| Correlation coefficient (mask)[b] | - | 0.84 | 0.86 |
| Bond length rmsd (Å) | - | 0.004 | 0.003 |
| Bond angle rmsd (°) | - | 0.625 | 0.539 |
| All-atom clashscore | - | 8.82 | 4.75 |
| Ramachandran plot | - | | |
| Outliers (%) | | 0.00 | 0.00 |
| Allowed (%) | | 2.49 | 1.60 |
| Favored (%) | | 97.51 | 98.40 |
| Rama-Z (whole) | - | -0.85 | 0.03 |

| | | | |
|---|---|---|---|
| CaBLAM Outliers (%) | | 0.87 | 0.56 |
| Rotamer outliers (%) | | 0.00 | 0.56 |
| PDB ID | - | 9FCI | 9FCJ |

[a]1% and 99% quantiles from local resolution computed in cryoSPARC with Adaptive Window Factor of 20

[b]Calculated in phenix

**Table S2.** Protein purification buffers.

| | Purification step (column)/Experiment | Buffer composition |
|---|---|---|
| USP1 or UAF1 | Lysis | 50 mM Tris pH 8.0, 150 mM NaCl, 5% (v/v) glycerol, 10 mM β-mercaptoethanol, 10 mM Imidazole, 2 mM $MgCl_2$, 1x cOmplete EDTA-free protease inhibitor cocktail, >10 units/mL benzonase |
| | Ni-NTA Wash 1/Subtractive | 50 mM Tris pH 8.0, 500 mM NaCl, 5% (v/v) glycerol, 10 mM β-mercaptoethanol, 10 mM Imidazole |
| | Ni-NTA Wash 2 | 50 mM Tris pH 8.0, 100 mM NaCl, 5% (v/v) glycerol, 1 mM TCEP, 10 mM Imidazole |
| | Ni-NTA Elution | 50 mM Tris pH 8.0, 75 mM NaCl, 5% (v/v) glycerol, 1 mM TCEP, 250 mM Imidazole |
| | Anion Exchange (ResourceQ 1 mL) | 50 mM Tris pH 8.0, 5% (v/v) glycerol, 1 mM TCEP, 100-1000 mM NaCl |
| | Gel Filtration (Superdex 200 Increase 10/300 GL) | 20 mM Tris pH 8.0, 150 mM NaCl, 5% (v/v) glycerol, 5 mM DTT |
| FANCD2, FANCI | Lysis | 50 mM Tris pH 8.0, 400 mM NaCl, 5% (v/v) glycerol, 5 mM β-mercaptoethanol, 10 mM Imidazole, 2 mM $MgCl_2$, 1x cOmplete EDTA-free protease inhibitor cocktail, >10 units/mL benzonase |
| | Ni-NTA Wash 1 | 50 mM Tris pH 8.0, 400 mM NaCl, 5% (v/v) glycerol, 5 mM β-mercaptoethanol, 10 mM Imidazole |
| | Ni-NTA Wash 2 | 50 mM Tris pH 8.0, 150 mM NaCl, 5% (v/v) glycerol, 1 mM TCEP, 10 mM Imidazole |
| | Ni-NTA Elution | 50 mM Tris pH 8.0, 100 mM NaCl, 5% (v/v) glycerol, 1 mM TCEP, 250 mM Imidazole |

| | | |
|---|---|---|
| | Anion Exchange (HP Q 5 mL) | 50 mM Tris pH 8.0, 5% (v/v) glycerol, 1 mM TCEP, 100-1000 mM NaCl |
| | Gel Filtration (Superose 6 Increase 10/300 GL) | 20 mM Tris pH 8.0, 400 mM NaCl, 5% (v/v) glycerol, 5 mM DTT |
| USP7$^{CD}$ | Lysis Buffer | 50 mM Tris pH 8.0, 150 mM NaCl, 5% (v/v) glycerol, 10 mM imidazole, 10 mM β-ME, 1 mM MgCl2, 1x cOmplete EDTA-free protease inhibitor cocktail, >10 units/mL benzonase |
| | Wash Buffer 1 | 50 mM Tris pH 8.0, 500 mM NaCl, 5% (v/v) glycerol, 10 mM imidazole, 10 mM β-ME |
| | Wash Buffer 2 | 50 mM Tris pH 8.0, 150 mM NaCl, 5% (v/v) glycerol, 10 mM imidazole, 10 mM β-ME |
| | Anion Exchange (ResourceQ 1 mL) | 20 mM Tris pH 8.0, 5% (v/v) glycerol, 10 μM β-ME, 100-1000 mM NaCl |
| | Gel Filtration (Superdex75 10/300 GL) | 20 mM Tris pH 8.0, 150 mM NaCl, 5% (v/v) glycerol, 5 mM DTT |
| Ubiquitin-prg | Lysis | 20 mM $Na_2HPO_4$ pH 7.2, 200 mM NaCl, 1 mM EDTA, protease inhibitor cocktail (30mL/1L cell culture) |
| | Wash Buffer 1 | 20 mM $Na_2HPO_4$ pH 7.2, 200 mM NaCl, 0.1 mM EDTA |
| | Wash Buffer 2 | 20 mM $Na_2HPO_4$ pH6.0, 200 mM NaCl, 1 mM EDTA |
| | Elution Buffer | 20 mM $Na_2HPO_4$ pH6.0, 200 mM NaCl, 1 mM EDTA, 200 mM MESNa |
| | Gel Filtration (Superdex75 16/600) | 1xPBS |
